# Use of massive DNA barcoding to monitor biodiversity: a test on forest soil macrofauna

**DOI:** 10.1101/2025.02.07.636998

**Authors:** Franck Jabot, Gwenaëlle Auger, Pauline Bonnal, Mathilde Pizaine, Marilyn Roncoroni, Sandrine Revaillot, Julien Pottier

**Affiliations:** INRAE, VetAgro Sup, UMR Ecosystème Prairial, Université Clermont Auvergne, Clermont-Ferrand, France

**Keywords:** Biodiversity, monitoring, DNA barcoding, soil fauna, megabarcoding

## Abstract

Biodiversity monitoring primarily focuses on particular taxonomic groups for which identification expertise is widely available, such as vascular plants or birds. As a result, the response of many taxonomic groups to forest management is much less documented. With the advent of low-cost next generation barcoding approaches, it is now possible to envisage monitoring strategies based on molecular approaches, beyond metabarcoding/eDNA strategies that do not give access to species abundances and integrate species presence over largely unknown spatio-temporal scales. In this contribution, we demonstrate the use of massive DNA barcoding – also referred as megabarcoding – as a promising solution to overcome identification difficulties in demanding taxonomic groups. We performed a proof of concept study in a mountainous beech forest in the Massif Central, France. We sampled soil macrofauna at 25 sampling sites and managed to visually identify at the species level 130 out of the 1413 sampled individuals. Using megabarcoding on the 1283 remaining non-identified individuals, we managed to assign 1124 additional individuals to an operational taxonomic species at a competitive cost. We present the summary statistics of barcoding success in the different taxonomic groups encountered and in larvae versus adult individuals. We demonstrate that larvae individuals, which can hardly be visually identified at the species level, make a substantial contribution to overall macrofauna diversity. We finally showcase how this megabarcoding approach provides a concrete avenue for forest biodiversity monitoring, by assessing the cost and labour intensity of this approach.

## 1. Introduction

Forest biodiversity monitoring mainly focuses on a limited number of taxonomic groups, among which plants, birds and beetles have attracted much attention (Paillet et al. 2010). Rationale for monitoring such particular taxonomic groups have been abundantly developed: these groups may harbour some ecological and/or conservation importance; they can potentially be used as indicator species; and their monitoring is practical, or at least feasible. Indeed, plants constitute habitats and resources for a wide-range of often specialized phytophagous invertebrates (Jaenike 1990), hence monitoring plants is also a way to indirectly monitor the potential presence of a series of interacting organisms (Basset et al. 2012, Castagneyrol and Jactel 2012). Plants can also be used as indicator species of a wide range of environmental (Tichy et al. 2023) and historic (Hermy et al. 1999) variables. Birds have also been proposed as potential indicator species of forest disturbance (Canterbury et al. 2000), as well as wood-inhabiting beetles (Nilsson et al. 2001). Wood-living beetles in particular are important components of forest biodiversity and play a key role in wood decomposition and nutrient cycling (Stokland et al. 2012). The functional composition of this taxonomic group is further influenced by forest management practices (Gossner et al. 2013) and is thus a group of conservation concern.

Focusing forest biodiversity monitoring on particular taxonomic groups also comes with a number of shortcomings. Indicator species remain surrogates of variables of interest and are also often influenced by the environmental, historical and landscape contexts in which they are applied (Gossner et al. 2014, Lalechère et al. 2017). Besides, several taxonomic groups may respond differently to human management or natural perturbations, rendering the generalization from one group to the others risky (Barlow et al. 2007, Cours et al. 2023). It is thus important to expand forest biodiversity monitoring to a larger number of taxonomic groups than what is currently performed (Nilsson et al. 2001, Burrascano et al. 2023). Among the often-neglected taxa, soil organisms account for ca. 60% of the total number of species on Earth (Anthony et al. 2023) and have an essential role in ecosystem functioning (Bardgett and Van der Putten 2014). Yet, relatively few monitoring programs focus on forest soil biodiversity (Korboulewsky et al. 2016).

A major obstacle to the study of the diversity of soil organisms is the rarity of suitable taxonomic expertise (Eisenhauer et al. 2017), and the difficulty or impossibility to identify larval stages that can substantially contribute to soil fauna communities (Paquin and Coderre 1997). Molecular approaches have been developed for more than two decades for taxonomic identification of a wide range of taxa (Hebert et al. 2003), including soil organisms (Rougerie et al. 2009, Decaens et al. 2013). They consist in sequencing a small region of the genome of the studied organism that is called the DNA barcode. The cost of DNA barcoding was however prohibitive until recently, limiting its widespread use in biodiversity monitoring programs. But this cost has dramatically decreased recently, thanks to the use of nanopore sequencing technologies and of low-cost protocols for DNA extraction and amplification (Srivathsan et al. 2021). In the meantime, reference libraries are continuously completed for a wide range of taxa (e.g., Rougerie et al. 2015, Santos-Perdomo et al. 2024). For instance, the Barcode of Life Database (BOLD) now gathers more than 19 million barcode sequences, and coordinated efforts to fill the remaining gaps are underway (Roslin et al. 2022). The massive barcoding of individual specimens therefore constitutes a promising avenue to speed-up and largely develop biodiversity monitoring, in particular for less studied taxa.

Barcoding a large number of individual specimens simultaneously has been recently coined as “megabarcoding” (Chua et al. 2023) to distinguish it from the “metabarcoding” approach in which mixtures of organisms or their living media (environmental DNA) are sequenced. The potential of metabarcoding, and in particular environmental DNA, for biodiversity monitoring has been abundantly developed in the literature (Rodriguez-Ezpeleta et al. 2021). The great advantage of metabarcoding is its speed of use: an environmental sample is quickly processed and its metabarcoding informs on a list of species that have left DNA traces in the sample. The main limitation of this approach is that the inference of species abundances from metabarcoding results remains challenging (Elbrecht and Leese 2015, Beng and Corlett 2020). Another limitation of the environmental DNA approach is that it integrates the DNA signal over spatio-temporal scales that are still difficult to appraise (Bloor et al. 2021). In contrast, megabarcoding enables to assess species abundances by processing individual specimens and to control the spatiotemporal scale of sampling, as traditional inventory methods. In particular, it should enable to track spatial and temporal changes in species relative abundances, which usually prelude species turnovers.

Given the rapid and promising developments of low-cost megabarcoding, comparisons of its efficiency among a full range of taxa composing each ecosystem compartments is now needed. Low-cost megabarcoding has been successfully applied to Dipteran specimens (Srivathsan et al. 2021, Hartop et al. 2024). A recent work using a higher cost protocol extended this application to a wider range of taxa, but mainly epigeal ones (Hebert et al. 2025). Soil macrofauna gathers various taxonomic groups with varied physiological and anatomical attributes that may compromise the ability of low-cost protocols to efficiently extract and amplify DNA, such as the presence of protective shells (in Styllommatophora), of dense exoskeletal cuticles (in Insecta, Isopoda and Diplopoda) or the presence of mucus (in Styllommatophora and Haplotaxida). For instance, mollusc slime has been shown to compromise amplification success (Chakraborty et al. 2020). Our aim in this study was thus to test the applicability of low-cost megabarcoding to monitor soil fauna communities in a temperate forest. We specifically assessed 1) whether the low-cost protocol proposed by Srivathsan et al. (2021) for the extraction, amplification and purification of DNA can be effectively adapted to soil fauna taxa, 2) the completeness of the BOLD database for the taxa encountered in our survey, and 3) the relative ability to monitor larval stages with DNA barcoding, compared to the one obtained for adult individuals.

## 2. Material and methods

### 2.1 Soil fauna collection and barcoding protocol

We performed an inventory of soil macrofauna in a beech forest stand located in Laqueuille in the French Massif Central (exact location: 45.6236 N, 2.7393 E, altitude: 1182m) in June 2022. This stand was rather homogeneous with low micro-habitat diversity, except some patches of *Vaccinum myrtillus* near the stand edge. Twenty-five soil samples were gathered within a 50 m square. Each sample consisted in hand sorted individuals larger than 3 mm extracted from soil monoliths of 18 cm x 18 cm and 10 cm deep together with the litter above. These individuals were then immediately stored in 96° ethanol. Individuals were first visually identified using a stereomicroscope (Zeiss Stemi 2000-C). This visual identification enabled us to sort the 1413 individuals into morpho-species that had variable taxonomic resolutions. Among them, 130 individuals were either identified at the species level, or were singletons of a morpho-species and were thus treated as a distinct species in the following. The remaining 1283 partially identified individuals were barcoded following the protocol detailed in Srivathsan et al. (2021).

This barcoding protocol consisted in rinsing each individual with purified water (Elga Labwater Purelab Ultra MK2) and placing it in one well of a 96-well PCR microplate. If the individual was too large to fit into a well, one of its legs (or anus for earthworms) was detached and put into the well. DNA extraction was performed using a HotShot solution (Truett et al. 2000) with a heating at 65°C for 18 minutes and 98°C for 2 minutes. Extracted DNA was then amplified within a mixture of 3 μL of extracted DNA, 7 μL of 2x Mastermix (CWBio CW0682L), 1 μL of BSA, 2 μL of tagged primers. Left and right primers were tagged with 9 bp tags developed by Srivathsan et al. (2024) so that each well had a unique combination of left and right tags. We used the BF3-BR2 primer set (Elbrecht and Leese 2017) to amplify a 418 bp region of the COI barcode for 808 individuals and the LCO1490-HCO2198 primer set (Folmer et al. 1994) to amplify a 658 bp region of the COI barcode for the remaining 475 individuals. We used these two different sets to compare their efficiency to amplify the COI “Folmer” region of soil fauna. A previous benchmark on 21 primer sets by Elbrecht et al. (2019) revealed that the BF3-BR2 primer set performed better than the classical LCO1490-HCO2198 set when analysing a Malaise trap sample. We wanted to test whether this enhanced performance was also recovered on soil taxa. We used the following PCR settings: 15 min at 95°C followed by 25 cycles of 30 s at 94°C, 1 min 30 s at 50°C and 1 min 30 s at 72 °C, followed by 10 min at 72°C. Amplified DNA was then pooled (fixed volume of 5 μL for each well) and a subset of 500 μL of this pooled DNA was purified using Ampure XP beads at 0.5 X. Library preparation for sequencing on a flongle followed the ONT protocol (with the modifications explained in Srivathsan et al. 2021). Srivathsan et al. (2021) recommended to pool up to 3 PCR plates on a flongle, while Hebert et al. (2025) demonstrated that up to 20 PCR plates could be pooled on the same device. We thus used an intermediate number and pooled between 6 and 10 PCR plates on each flongle. Note that to decrease barcoding costs, PCR success was not assessed with gel electrophoresis.

### 2.2 Bioinformatic processing

Raw sequences were then processed with the software ONTbarcoder 2.0 (Srivathsan et al. 2024) and only quality-checked consensus barcodes (i.e., no ambiguities nor insertion/deletion) for each well were retained. The individuals for which the first sequencing round failed were re-processed from scratch (extraction, amplification, purification and library preparation) for a second round of sequencing, and those still in failure were processed a third time. The barcodes obtained were compared to the BOLD database (Ratnasingham and Hebert 2007) using Boldigger3 (Buchner and Leese 2020). We used the animal library (public and private), the rapid search option, and similarity thresholds of 99%, 97% and 90% for species, genus and family respectively. This step was performed on the twenty first of May 2025. This choice of using this user-friendly identification engine was motivated to keep the process simple and easy to reproduce, although it comes with a potential cost of neglecting some reference sequences that may be present in other databases, notably GenBank. We then clustered barcodes that could not be matched to a species on the BOLD database into distinct OTUs (operational taxonomic units) using a UPGMA method and a similarity threshold of 99% (Buchner and Leese 2020). We replicated our analyses using alternative thresholds of 98% and 97% to assess the sensitivity of our results to this arbitrary threshold and obtained very similar results (Figure S9).

### 2.3 Data analysis

We computed the barcoding efficiency as the percentage of consensus barcodes obtained (the number of obtained barcodes divided by the number of sequenced individuals). We repeated this computation for each of the three rounds of sequencing and separately for this list of taxonomic groups: Arachnida, Chilopoda, Diplopoda, Diplura, Haplotaxida, Insecta, Isopoda and Stylommatophora.

We also computed the barcoding efficiency for each set of primers (BF3/BR2 and LCO1490/HC02198) separately. To test whether the BF3-BR2 primer set lead to a larger barcoding efficiency than the LCO1490-HCO2198 set, we fitted a Bernouilli GLM explaining barcoding success of an individual as a function of the primer set used and of the taxonomic group of the individual. This analysis was restricted on data of the first round of barcoding and was replicated with and without interaction terms between primer set and taxonomic groups.

We computed OTUs accumulation curves to assess the completeness of our species inventory, distinguishing the adult-only and total assemblages of soil fauna individuals. This was performed with the function “specaccum” of the vegan R package (Oksanen et al. 2007). We finally compared the summary statistics of inventories based either on visual (morphospecies) or on barcoding-based identification by computing species/OTU abundance distributions, richness, Shannon’s index, Pielou’s equitability index and average abundance-based Bray-Curtis dissimilarity between samples.

Datasets to replicate the analyses are included in the Supplementary Information (Tables S1 and S2) as well as the corresponding R script (Text S3). This material is also available on public repositories (urls can be found at the end of the manuscript).

## 3 Results

Using a low-cost DNA barcoding protocol (see Tables S4 and S5 for detailed costs), we obtained an overall barcoding efficiency of 88 %: we recovered 1124 DNA barcodes out of the 1283 individuals processed (Fig. 1A). These barcodes clustered into 122 distinct Operational Taxonomic Units (OTUs) among which 65 (53 %) had a species name in the BOLD database (Fig. 1B). Interestingly, OTUs without species names in the BOLD database were not solely the rarest taxa in our samples (Fig. 1C). We obtained a larger barcoding efficiency on larvae (96 % of the 234 larvae processed) than on adult individuals (86 % of the 1049 adults processed). Barcodes of larvae had moreover a larger percentage of species name presence in the BOLD database (77 % versus 37 %).

**Figure 1:**
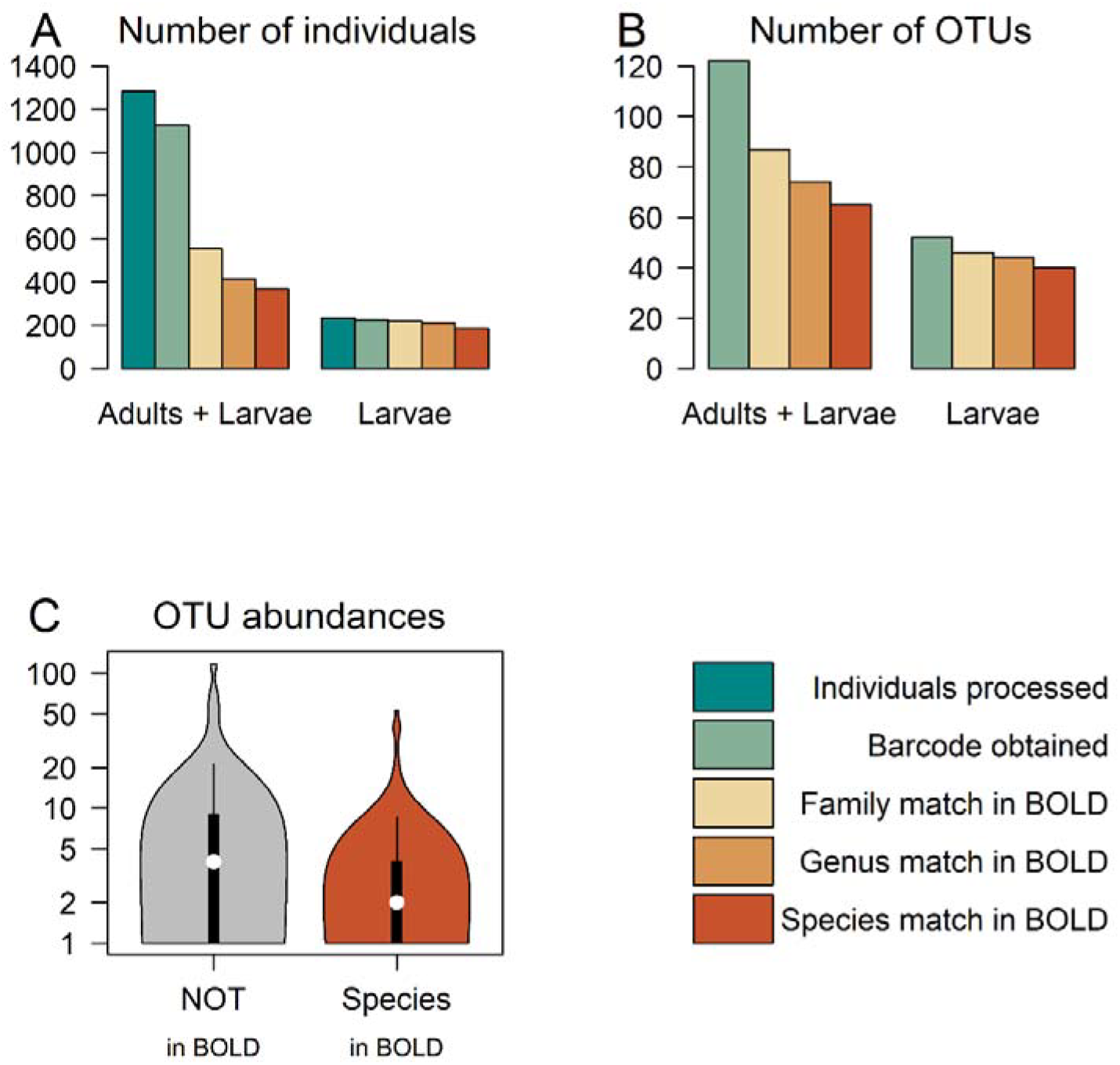
DNA barcoding efficiency of soil macrofauna. Panel A: number of individuals for which DNA barcodes have been obtained and the corresponding matches at various taxonomic levels in the BOLD database. Panel B: number of Operational Taxonomic Units (OTUs) and the corresponding matches at various taxonomic levels in the BOLD database. Panel C: Distribution of OTU abundances among OTU with a species name present or absent in the BOLD database.

DNA barcoding efficiency was highly heterogeneous among taxonomic groups (Figure 2), varying between 22 % (for Stylommatophora) and 100 % (for Arachnida). The percentage of species name presence in the BOLD database was also highly heterogeneous among these groups, varying between 0.3 % (for Diplura) and 82 % (for Insecta). We observed some variation in these numbers depending on the primer set used; the barcoding success among taxonomic groups for each primer set separately is depicted in Supplementary Figure S8.

**Figure 2:**
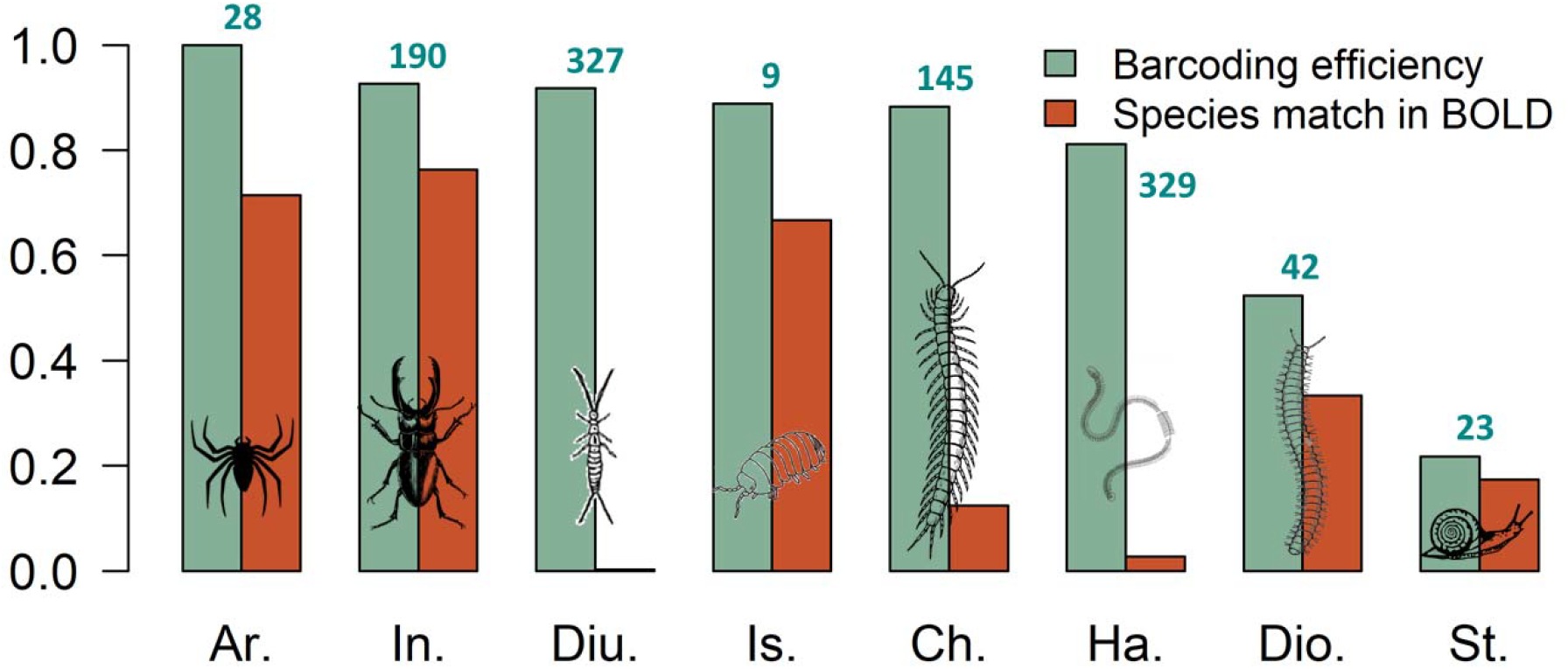
Heterogeneity of DNA barcoding efficiency among taxonomic groups. Barcoding efficiency is equal to the proportion of individuals in each taxonomic group for which a DNA barcode could be obtained. Species match is the proportion of individuals for which a species match was obtained in the BOLD database. Ar: Arachnida, In: Insecta, Diu: Diplura, Is: Isopoda, Ch: Chilopoda, Ha: Haplotaxida, Dio: Diplopoda, St: Stylommatophora. The numbers above the bars correspond to the number of individuals processed in each taxonomic group.

Our study also revealed that DNA barcoding failure could largely be reduced by re-processing failed individuals (Figure 3). Using this approach, barcoding efficiency sequentially improved from 59 % (using a single barcoding round for all individuals), to 81 % (using a second barcoding round for individuals in failure) up to 88 % (using a third barcoding round for individuals in failure after the first two rounds). The barcoding efficiency of the first two rounds was close (59 % and 56 % for round 1 and 2 respectively), while the one of the third round was lower (36 %). Finally, we observed a sizeable difference in barcoding efficiency among the two pairs of primers used (68 % and 43 % for BF3/BR2 and LCO1490/HC02198 respectively). This difference was significant (*p*<0.001) and did not interact significantly with taxonomic groups (see Supplementary Text S7 for the full results).

**Figure 3:**
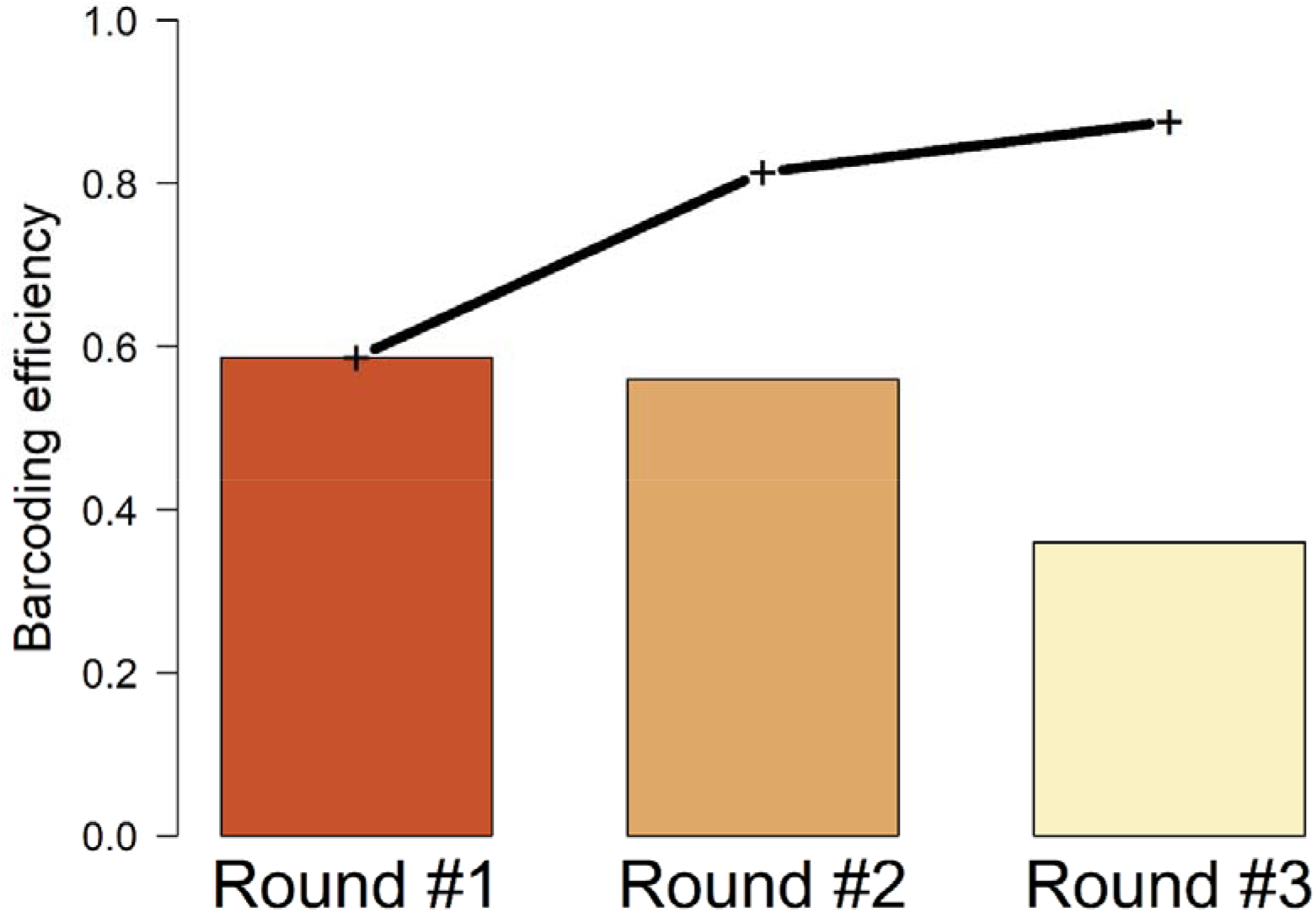
Improvement in DNA barcode efficiency brought by multiple rounds of barcoding. Each bar height stands for the barcoding efficiency of the barcoding round. The broken line represents the cumulative barcoding efficiency after a given number of barcoding rounds.

Our study further revealed that 25 samples of soil monoliths of 18 cm x 18 cm x 10 cm were not sufficient to reach an asymptote in the OTUs accumulation curve (Figure 4). Interestingly, OTUs accumulated faster when including larvae individuals, although larvae only accounted for 18 % of the total number of individuals. This was linked to the fact that 39 of the 148 OTUs (26 %) were only found among larvae individuals.

**Figure 4:**
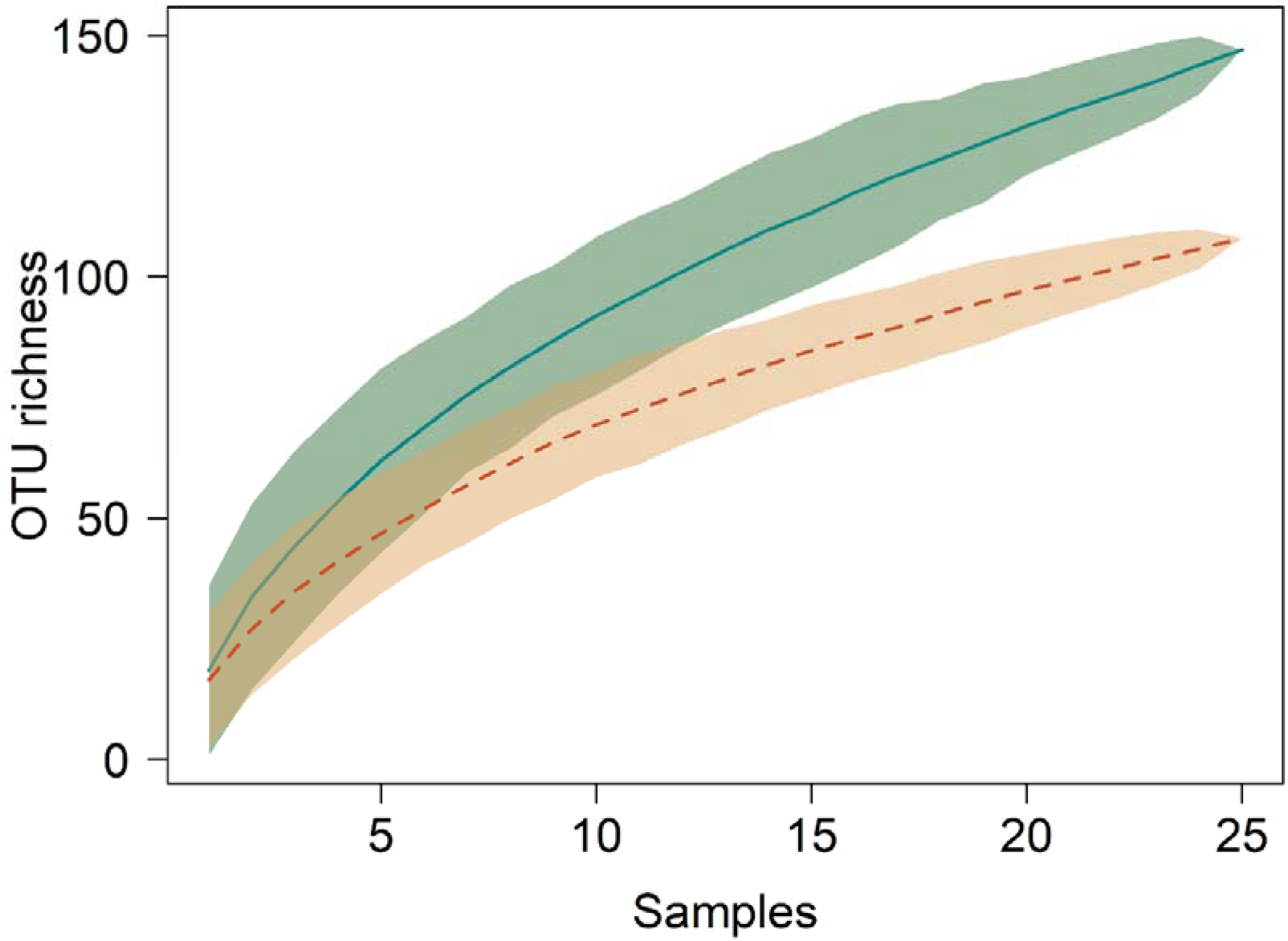
OTU accumulation curves as a function of the number of samples considered. The plain blue curve represents the OTU accumulation curve for the total community (adult and larvae individuals) while the dashed orange curve represents the OTU accumulation curve for the adult individuals only. Light colour areas depict confidence intervals.

Our analysis finally revealed that the barcoding-based species inventory strongly modified the initial morphospecies-based inventory, by more than doubling the number of OTUs detected (from 74 morphospecies to 148 OTUs, Figure 5B) and strongly altering biodiversity indices such as Shannon’s index (Fig. 5C), Pielou’s equitability index (Fig. 5D) and average Bray-Curtis dissimilarity between samples (Fig. 5E). The barcoding-based OTU abundance distribution was consequently strongly altered by being much more even than the original morphospecies-based abundance distribution (Figure 5A). The taxonomic resolution of the barcoding-based inventory was also sharply improved with the number of individuals with a species-level identification multiplied by more than three, from 130 individuals to 488 individuals.

**Figure 5:**
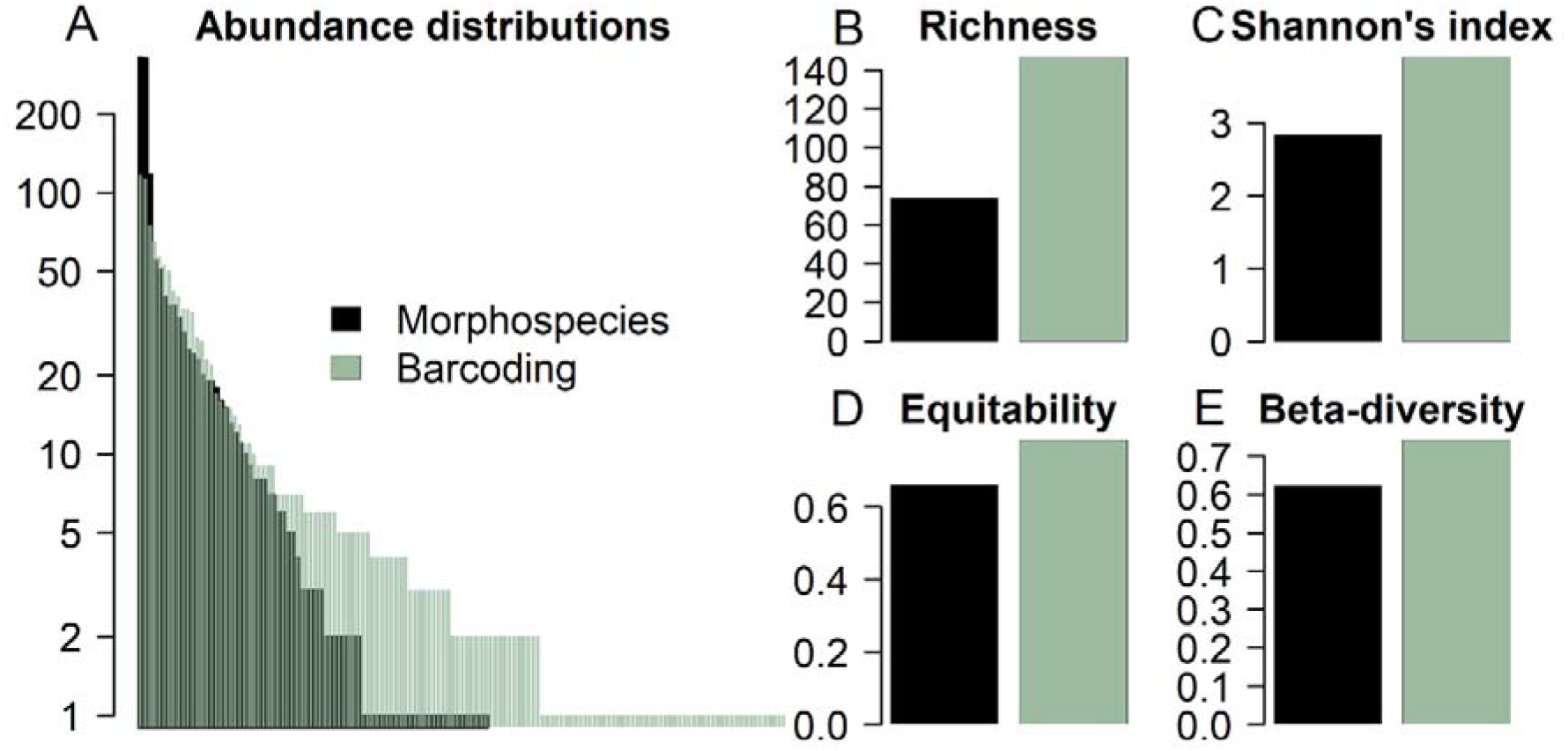
Comparison of the morphospecies-based and the barcoding-based abundance distributions (A), richness (B), Shannon’s index (C), Pielou’s equitability index (D) and average abundance-based Bray-Curtis dissimilarity index between samples (E).

## 4. Discussion

Our study demonstrates a clear success of megabarcoding to assess the diversity of forest soil macrofauna. Nearly 90% of the processed individuals were successfully barcoded and we were able to attribute a species name to nearly half of them (Fig. 1). Since taxonomic expertise of soil organisms is particularly rare, megabarcoding of sampled individuals could therefore constitute an operational way to perform soil biodiversity monitoring. Another major advantage of megabarcoding of soil macrofauna is the identification of larvae. In our study case, we found that larvae accounted for 18% of sampled individuals, and contained a large number of OTUs not found among adult individuals (26% of the total OTU number). Therefore, the inclusion of soil larvae in biodiversity assessment, enabled by megabarcoding, may profoundly alter our understanding of the rich forest soil fauna diversity (Fig. 4 and 5).

Our study further reveals that barcoding efficiency is variable among taxonomic groups, as previously demonstrated by Hebert et al. (2025). Despite this variation, barcoding efficiency was good to very good (above 80 % or 90 %) for most taxonomic groups, except for Stylommatophora and to a lesser extent Diplopoda (Fig. 2). This may explain why barcoding efficiency was remarkably large (96.1 %) for larvae individuals that mostly belonged to Coleoptera, Araneae and Diptera and thus not to these more problematic taxonomic groups (Table S1). The lower barcoding efficiency for the Stylommatophora and Diplopoda individuals echoes previous studies that reported barcoding difficulties in these taxonomic groups (Spelda et al. 2011, Chakraborty et al. 2020). It may be due to a variety of reasons (notably, failure of or poor DNA extraction, lack of or low DNA amplification) that would need to be further investigated. Note that Stylommatophora individuals were not extracted from their shells before DNA extraction. This may be part of the explanation for the lower barcoding efficiency of this group, on top of mucus-related amplification difficulties (Chakraborty et al. 2020). The number of processed individuals for these two groups was nevertheless low (23 and 42 respectively, Fig. 2) so that additional tests will be required to confirm or not this lower barcoding efficiency. And solutions to this problem may be sought, for instance by lengthening the heat extraction of DNA in the HotShot solution, by testing alternative primer sets for these groups (e.g., “C_LepFolF-C_LepFolR” as in Spelda et al. 2011), or by sequencing individuals of these groups separately from the other taxonomic groups. This last solution may decrease the among-individual variability in numbers of sequencing reads, thereby increasing the number of reads obtained for the individuals of these difficult groups.

The barcoding efficiency that we reached on the first two rounds was slightly above 50 % (Fig. 3). This is lower than the ones reported by previous studies that were mostly in the range of 80-90 % (Srivathsan et al. 2024, Hebert et al. 2025). Various factors can be put forward to explain the lower barcoding efficiency that we obtained in this study. A first explanation lays in the taxonomic groups studied. Previous megabarcoding studies have notably targeted Dipteran individuals (Srivathsan et al. 2021) that lead to larger barcoding efficiencies than Coleopteran and Arachnid individuals (Hebert et al. 2025). A second explanation linked to the first one is that we pooled individuals of different taxonomic groups in the same flongle. Sorting individuals of different taxonomic groups and sequencing them separately may enable some enhancement in barcoding efficiency, since the DNA sequences of more productive taxonomic groups would not overwhelm the ones of less producing groups. A third explanation lays in the number of microplates that we pooled on each flongle for sequencing. Srivathsan et al. (2024) recommended to pool up to 3 microplates on a flongle, while Hebert et al. (2025) demonstrated that up to 20 microplates could be pooled on a single flongle, but with the use of high-fidelity Taq polymerases for DNA amplification. We chose intermediate numbers and pooled between six and ten microplates on each flongle. Although this strategy led to a smaller barcoding efficiency, we found that it was a good compromise to speed up the treatment of samples, since the library preparation steps are long relative to the extraction and amplification ones (Table S6), and to further decrease barcoding costs (Table S4). Importantly, by re-sequencing individuals in failure of barcode recovery, we demonstrated that final barcoding efficiency was comparable to the one of previous studies (Fig. 3). For this, we recommend using two rounds of barcoding, since the third round provides comparatively fewer barcodes (Fig. 3).

As argued before by Srivathsan et al. (2021), the cost of barcode sequencing has dramatically decreased thanks to the use of nanopore technologies. But the total cost of DNA barcoding also incorporates the costs of reagents involved in the extraction and amplification of DNA. The protocol of Srivathsan et al. (2021) put forward a number of innovations to cut down these costs, including the use of low-cost HotShot solutions to extract DNA instead of commercial kits and the use of low PCR reaction volumes. We provide in Table S4 the details of the reagent costs implied in the full protocol and compute that the production of a DNA barcode in this study costed around one euro. Although this cost depends on the commercial arrangements each lab may get with reagent suppliers, this nevertheless gives the order of magnitude of megabarcoding costs. These costs could be further decreased by using MinION flow cells instead of Flongles for larger sample sizes. We also detail the equipment costs to perform megabarcoding and demonstrate that it amounts to around 20,000 euros for minimal equipment needs (Table S5). Finally, following this protocol, a single lab technician can barcode around 1000 (2000) individuals per week if the lab is equipped with a single (two) thermocycler(s) and thus obtain around 500 (1000) barcodes (Table S6).

A current limitation of megabarcoding is that reference databases are still lacunar, especially for soil fauna (Recuero et al. 2024). In our study, nearly half of barcodes could not be matched to a referenced species in the BOLD database. This limitation is however not definitive. Indeed, reference databases are updated at a rapid pace: the number of sequences in the BOLD database has increased by 67 % in 2024. Besides, coordinated regional and national efforts are under way to barcode missing taxa (e.g., Spelda et al. 2011, Hendrich et al. 2015, Roslin et al. 2022). This implies that currently unmatched sequences could be matched in the future, so that inventories based on megabarcoding can be progressively updated, a feature especially important for longitudinal and/or comparative studies. To illustrate this point, during the time interval of the review process, the number of individuals that could be matched to a species in the BOLD database increased by 10% (i.e., 44 additional individuals).

## 5. Conclusion

We demonstrated that the low-cost megabarcoding protocol of Srivathsan et al. (2024) could be fruitfully applied to forest soil macrofauna and that it led to an overall very good barcoding efficiency of nearly 90 % (Fig. 1). Two taxonomic groups, Styllommatophora and Diplopoda, would require further study to reach such high level of barcoding efficiency (Fig. 2). Megabarcoding of soil fauna also enables the identification of larvae individuals. Our study demonstrates that these larvae individuals substantially contribute to macrofauna biodiversity (Fig. 4), with many larvae OTUs not encountered among adult individuals. Finally, megabarcoding enables to keep track of OTUs abundances and therefore provides a strongly renewed assessment of soil macrofauna community structure (Fig. 5). With a cost of ca. one euro per barcode, megabarcoding constitutes a concrete avenue to vastly develop the monitoring of forest soil macrofauna.

## CRediT authorship contribution statement

**Franck Jabot:** Conceptualization, Data curation, Formal analysis, Funding acquisition, Investigation, Methodology, Project administration, Supervision, Writing – original draft. **Gwenaëlle Auger**: Data curation, Investigation, Writing – review and editing. **Pauline Bonnal**: Data curation, Methodology, Investigation, Writing – review and editing. **Mathilde Pizaine**: Data curation, Investigation, Writing – review and editing. **Marilyn Roncoroni**: Investigation, Writing – review and editing. **Julien Pottier**: Conceptualization, Supervision, Writing – review and editing.

## Data availability statement

Datasets to replicate the analyses are included in the Supplementary Information (Tables S1 and S2) as well as the corresponding R script (Text S3). This material has also been deposited on public repositories, the datasets can be found at doi:10.57745/LPYOC1 and the R script at Link2.

## Acknowledgements

This work was supported by the French Agence Nationale de la Recherche [grant number ANR-20-CE32-0004], by the IDEX-ISITE initiative [grant number 16-IDEX-0001(CAP 20-25)] and by Clermont Auvergne Metropole. We also thank Amrita Srivathsan for her advices to replicate their low-cost barcoding protocol and Isabelle Bosio for her help during field work.

## Supplementary Information

Table S1: Barcoding-enhanced inventory data. This dataset includes the identification of 1124 forest soil macrofauna individuals by DNA barcoding and the visual identification of 130 additional individuals.

Table S2: Visual inventory data. This dataset includes the visual identification of 1413 forest soil macrofauna individuals.

Text S3: R script to replicate the analyses and figures.

Table S4: Data on reagent costs.

Table S5: Data on equipment costs.

Table S6: Duration of each barcoding step.

Supplementary Text S7: Full results of the statistical analysis.

Supplementary Figure S8: Barcoding success among taxonomic groups for each primer set.

Supplementary Figure S9: Summary statistics of macrofauna biodiversity for different similarity thresholds to delineate OTUs.

